# Viral capsid, antibody, and receptor interactions: experimental analysis of the antibody escape evolution of canine parvovirus

**DOI:** 10.1101/2023.01.18.524668

**Authors:** Robert A. López-Astacio, Oluwafemi F. Adu, Daniel J. Goetschius, Hyunwook Lee, Wendy S. Weichert, Brian R. Wasik, Simon P. Frueh, Brynn K. Alford, Ian E.H. Voorhees, Joseph F. Flint, Sarah Saddoris, Laura B. Goodman, Edward C. Holmes, Susan L. Hafenstein, Colin R. Parrish

## Abstract

Canine parvovirus (CPV) is a small non-enveloped single-stranded DNA virus that causes serious diseases in dogs worldwide. The original strain of the virus (CPV-2) emerged in dogs during the late-1970s due to a host range switch of a virus similar to the feline panleukopenia virus (FPV) that infected another host. The virus that emerged in dogs had altered capsid receptor- and antibody-binding sites, with some changes affecting both functions. Further receptor and antibody binding changes arose when the virus became better adapted to dogs or to other hosts. Here, we use *in vitro* selection and deep sequencing to reveal how two antibodies with known interactions select for escape mutations in CPV. The antibodies bind two distinct epitopes, and one largely overlaps the host receptor binding site. We also engineered antibody variants with altered binding structures. Viruses were passaged with the wild type or mutated antibodies, and their genomes deep sequenced during the selective process. A small number of mutations were detected only within the capsid protein gene during the first few passages of selection, and most sites remained polymorphic or were slow to go to fixation. Mutations arose both within and outside the antibody binding footprints on the capsids, and all avoided the TfR-binding footprint. Many selected mutations matched those that have arisen in the natural evolution of the virus. The patterns observed reveal the mechanisms by which these variants have been selected in nature and provide a better understanding of the interactions between antibody and receptor selections.

**IMPORTANCE:** Antibodies protect animals against infection by many different viruses and other pathogens, and we are gaining new information about the epitopes that induce antibody responses against viruses and the structures of the bound antibodies. However, less is known about the processes of antibody selection and antigenic escape and the constraints that apply in this system. Here, we use an *in vitro* model system and deep genome sequencing to reveal the mutations that arise in the virus genome during selection by each of two monoclonal antibodies or their engineered variants. High-resolution structures of each of the Fab: capsid complexes revealed their binding interactions. The engineered forms of the wild-type antibodies or mutant forms allowed us to examine how changes in antibody structure influence the mutational selection patterns seen in the virus. The results shed light on the processes of antibody binding, neutralization escape, and receptor binding, and likely have parallels for many other viruses.

## INTRODUCTION

Antibody binding and neutralization, along with antigenic variation, are fundamental properties of the interactions between viruses and vertebrate hosts (1, 2). While many features of the interactions between viruses and their host antibody responses are known, there are several aspects that remain to be determined, or that appear to vary depending on the viruses or hosts involved. These include the nature of the antigenic structures of the viruses, the development and genetic diversity of host antibody responses, the nature and selection of antibody escape mutations, and clarifying how those interact with essential viral functions such as receptor binding sites (3–6). Here we investigate the experimental evolution of parvoviruses in the face of binding of antibodies with known structural interactions.

Canine parvovirus (CPV) (carnivore protoparvovirus-1) emerged as a new pathogen in dogs in the late-1970s, and it is an evolutionary variant of a virus similar to the feline panleukopenia virus (FPV) of cats (7). The first strain of CPV that spread worldwide in 1978 was designated CPV type-2 (CPV-2) to distinguish it from the previously described minute virus of canines (canine bocaparvovirus-1) (8). However, that original CPV-2 strain was replaced worldwide between 1979 and 1980 by a canine host-adapted and antigenic variant that had several sequence changes, mainly in the capsid protein gene, and that variant was designated CPV-2a (9). The CPV-2a sequence is the common ancestor of all CPV strains found in dogs after about 1980, which have continued to circulate among dogs globally and have also infected or spread among several hosts of the order Carnivora, including raccoons (7, 9–11).

The parvovirus capsid is a T=1 icosahedron assembled from a total of 60 copies of a combination of the viral capsid proteins (VPs), including 5 to 10 copies of VP1 and 55 to 50 copies of VP2. The capsid has several exposed features, including a raised region around the threefold axis (the threefold spike), a cylinder made up of five flexible loops that surround a pore that penetrates through each fivefold axis, and a depressed region (dimple) that spans the twofold axis (12). The CPV or FPV capsid binds to a variety of host ligands, including the host transferrin receptor type-1 (TfR) (13, 14), sialic acid (specifically N-glycolyl neuraminic acid; Neu5Gc) (15), and antibodies (16, 17).The TfR binds through its apical domain to a small region on the three-fold spike of the capsid, while Neu5Gc binds to a site within the twofold dimple (15, 18). The CPV capsid is highly antigenic and soon after infection antibodies are produced that recognize specific epitopes on the surface of the virus, primarily structures composed of multiple loops of one or more VP2 subunits (19). These antibodies provide solid protection against reinfection to infected and recovered or vaccinated animals, and maternally transferred antibodies in the colostrum also protect young animals against infection for several weeks after birth (20, 21). Most antibodies neutralize the parvovirus when present as IgGs, but the neutralization by the Fabs derived from those IgGs may vary widely (1, 22). The TfR binding site on the capsid has been mapped using cryoEM analysis, and overlaps the binding sites of many antibodies (14, 23). Many natural or experimentally selected mutations in the virus simultaneously alter both receptor and antibody binding, where either host range variation arises after antibody selection, or antigenic variation arises after TfR selection (24–27). Although many of those linked changes result from the overlapping structures of the receptor and antibody binding sites, it is unclear how the different mutations arise and are selected during the evolution of the viruses (28).

Antibodies are a key component of adaptive immunity in vertebrates. After natural parvoviral infection or post-vaccination, B cells that are bound by viral antigens are activated, and the highest affinity B cells selected expand most efficiently, and both develop the sustained antibody response and establish the memory cell population (29–32). Mouse or rat monoclonal antibodies (mAb) prepared against FPV, CPV-2, or CPV-2a capsids recognize many positions on the capsid surface, and cryoEM analysis of the binding sites of eight different rodent IgGs showed that their footprints covered about 65% of the capsid surface (1, 17). Near-atomic resolution structures of some of these antibodies bound to CPV capsids have been obtained and confirm that both the antibody heavy and light chain complementarity determining regions (CDRs) contact multiple surface loops from different VP2 (or VP1) subunits (33, 34).

Antigenic variation in CPV- and FPV-related viruses have frequently been observed during their natural evolution (35–37), and escape mutations are readily selected by the growth of viruses in the presence of the mAbs (38, 39). These natural or experimentally selected escape mutations tend to be clustered into a small number of positions on the capsid surface (16, 17). The binding sites of many antibodies on the capsids overlap the attachment site of the TfR, and naturally emerging and experimentally selected mutations may influence the binding or functions of both ligands (25, 26).

Here, we studied two antibodies that bind to distinct positions on the CPV capsid, both of which we have high-resolution structures of their capsid-bound forms, and we also produced engineered versions of these antibodies. The wild-type or engineered antibodies were mixed with CPV that was recently derived from plasmid clones – and which therefore contained few pre-existing variants –, and after passage of the viruses in the presence of the antibodies deep sequencing allowed us to follow the emergence of mutations in the viral genome.

## MATERIALS AND METHODS

### Mammalian and insect cells

Norden Laboratories feline kidney (NLFK/ cat) cells were grown in 1:1 McCoy’s 5A-Leibovitz L15 medium (Corning Life Sciences) supplemented with 5% fetal calf serum (FCS) and incubated at 37ºC and 5% CO_2_. NLFK cells were used for transfections with WT and mutated CPV-2 plasmids (virus stocks preparations), as well as for the experimental passages under the selection of antibodies and competition assays. Sf9 (*Spodoptera frugiperda-9*; Invitrogen) and High Five (*Trichoplusia ni*; Boyce Thompson Institute at Cornell University) (40) insect cells were grown in Grace’s insect medium (Invitrogen) with 10% FCS and Express Five insect serum-free medium (Invitrogen) with 2.4% L-Glutamine and 10% FCS, respectively, at 28ºC. Both insect cells were used to produce scFv-Fcs in a baculovirus-based protein expression system.

### Infectious CPV-2 plasmids and wild-type virus stocks

The infectious WT CPV-2 plasmid (plasmid pBI265 for CPV-2 strain) was produced in DH10B cells and the plasmid purified from single-bacterial colonies using the E.Z.N.A.® Plasmid Mini Kit I (Omega Bio-tek, cat. no. D6942-01). NLFK cells were transfected using Lipofectamine 2000 (Thermo Fisher Scientific, cat no. 11668019) with purified WT CPV-2 plasmid as per the manufacturer’s instructions. The cells were then immunostained using a mouse mAb conjugated to Alexa-Fluor 488 (CE10 in a 1:500 dilution) against the viral protein NS1 to reveal the transfection efficiency and viral replication. Virus stocks were collected as cell culture medium 10 days post-transfection, clarifying at 4,000 RPM for 5 minutes at 4ºC, and stored at -80ºC. Viruses were detected in supernatants through hemagglutination assay (HA) with a 0.5% (v/v) suspension of feline erythrocytes (41, 42). Both the WT CPV-2 plasmid and the initial virus stock were deep-sequenced to determine the sequence and variation of the initial input virus (see below for more details).

### Single mutant viruses

Antibody-selected mutants I101T, G224E, and N426D (with non-synonymous changes of residues in the VP2 gene) were prepared as single mutants in a WT CPV-2 background (infectious plasmid (43)) using Phusion™ High-Fidelity DNA Polymerase PCR-based mutagenesis as per manufacturer instructions. Mutated plasmids were Sanger sequenced to confirm that only the desired mutation was present, and viral stocks were deep sequenced in the course of the experiments. Some single mutant plasmids did not produce viable viruses, and their mutated regions were recloned via Gibson assembly into the appropriate infectious plasmid background to ensure that no additional mutations were present and to replace the terminal palindromic sequences at the 3’- and 5-’ends required for viral replication. Transfection efficiency and active viral replication were assessed as described above. Viruses were detected in supernatants through HA (41, 42). Virus stocks for each mutant were prepared as described above.

### Soluble mouse mAb IgG production and purification

The mouse or rat mAb used have been described previously (44). CPV-2-specific mouse mAb IgG Fab14 and rat mAb IgG FabE were purified from hybridoma supernatants by fast protein liquid chromatography (FPLC) using a 5 mL protein-G affinity column and eluted at pH 3.5. The IgG-containing fractions were combined and buffer-exchanged using a Merck Amicon™ Ultra centrifugal filter tube (Merck™, cat no. 10581342) in sterile PBS. After concentration, each IgG was quantified using the Pierce™ BCA Protein Assay Kit as per the manufacturer’s recommendations (Thermo Fisher Scientific, cat. no. 23225).

### Soluble CPV-2-specific WT scFv-Fcs

To allow ready manipulation of the antibody sequences and structures, the heavy and light chain variable domains of the Fab14 and FabE were each synthesized, and those were linked by three repeats of Gly-Gly-Gly-Ser to form a single-chain variable fragment (scFv) (25). Each sequence was fused to a baculovirus gp68 secretory signal sequence at the N-terminus and linked through a flexible linker to the Fc portion of human IgG-1 with a 6-His tag at the C-terminus to form the complete single-chain variable fragment and crystallizable fragment (scFv-Fc). Those were cloned into the pFastBac™ vector (Thermo Fisher Scientific, cat no. 10360-014), and bacmids produced by recombination with DH10Bac™ vectors in *E. coli* (Thermo Fisher Scientific, cat no. 10359-016). Cloned recombinant bacmids were analyzed through PCR, and recombinant bacmids were purified using the E.Z.N.A.® Plasmid Mini Kit I (Omega Bio-tek, cat. no. D6942-01). Baculoviruses were produced by transfecting Sf9 cells with recombinant bacmids using TransIT^®^-Insect Transfection Reagent (Mirus, cat no. MIR 6100), grown for 7 days at 28ºC, then stocks prepared in the same cells. The Fab14 and FabE scFv-Fc proteins were produced in High Five (Hi5) cells. The supernatants were collected three days after infection and dialyzed to buffer exchange in sterile PBS. The scFv-Fcs were purified through their included 6His tag by FPLC using a 5mL Ni-NTA column and eluted with 250mM imidazole after washes with 25mM imidazole (both at pH 8.0). Each scFv-Fc was buffer-exchanged using a Merck™ Amicon™ Ultra Centrifugal Filter tube (Merck™, cat no. 10581342) in sterile PBS, quantified following the Pierce™ BCA Protein Assay Kit protocol (Thermo Fisher Scientific, cat. no. 23225) and tested through an enzyme-linked immunosorbent assay (ELISA) against CPV-2.

### Engineering the antigen-binding site of CPV-2-specific scFv-Fcs

Predicted antibody: capsid interactions were modeled on the cryoEM structures (33, 34), and mutations selected using the swappa interactive tool available in UCSF Chimera X (45) following the Dunbrack model (46). New charged amino acid residues introduced into the antigen-binding sites were predicted to form distinct interactions with residues expressed on the CPV-2 capsid loops (**Figs. 4B and C**). Single or double scFv-Fc mutants were generated by changing hydrophobic or uncharged residues to charged amino acids within the antigen-binding site of either the light chain (Lc) or heavy chain (Hc) of the Fab14 or FabE scFv-Fc. The scFv-Fc amino acid side chains modified were <6.5 Å from oppositely charged side chains on the viral capsid, and the models are shown in **Fig. 4B and C**. For Fab14, light chain residue Asn-30 was changed to an Asp - predicted to interact with VP2 His-222, and heavy chain residue Thr-30 was changed to a Lys - predicted to interact with VP2 Asp-88. For FabE scFv-Fc, heavy chain residue Met-152 was changed to Asp - predicted to interact with VP2 residue Lys-570, and heavy chain Pro-202 was changed to Glu - predicted to interact with VP2 residue His-234. The scFv-Fcs mutant proteins were purified and quantified as described above.

### Measurement of relative scFv-Fc binding to CPV-2 capsids using ELISA

All scFv-Fc samples were tested via ELISA as previously described (47). Briefly, 96-well Nunc MaxiSorp plates (Thermo Fisher Scientific) were coated overnight at 4°C with 3 μg/mL CPV-2 empty capsids in carbonate buffer (pH 9.6) and then blocked for 2 hrs at 37°C using 5% BSA in PBST. The scFv-Fc (WT and mutants) were added at an initial standardized concentration of 10 μg/mL and serially diluted and incubated at room temperature for 1 hr. Wells were then washed three times with PBS with 0.05% Tween-20, then incubated with horseradish peroxidase (HRPO) conjugated donkey anti-human IgG (Fcγ fragment specific) antibody (Jackson ImmunoResearch Inc., West Grove, PA), and developed using 3,3’,5,5’-tetramethylbenzidine (TMB/ Thermo Fisher Scientific). Optical density (OD) values were recorded at 450 nm with a Multiskan Ascent plate reader (Thermo Fisher Scientific).

### scFv-Fc binding analysis using biolayer interferometry

To examine the effects of antibody mutations on binding, we incubated CPV-2 capsids to wildtype or engineered scFv-Fc using biolayer interferometry (BLI) (BLITz ForteBio, Menlo Park, CA). Protein A biosensors were blocked and hydrated in kinetics buffer (PBS with 0.02% ovalbumin and 0.02% Tween-20) at room temperature, scFv-Fcs bound to a constant level, and then those were used to measure capsid binding. Binding studies included a 30 s incubation in buffer to establish a baseline, 120 s loading scFv-Fc (either WT or mutated) in kinetics buffer to a consistent level of 400 nm, 30 s wash in kinetic buffer, 120 s incubation with purified WT CPV-2 capsids (500 μg/ml of protein) in kinetics buffer, and 120 s wash in kinetics buffer.

### ELISA measurement of relative mAbs binding to mutated CPV-2 capsids

To test the effects of the antibody-selected mutations in CPV-2 capsids (VP2 residues I101T, G224E, or N426D) on binding to both mAbs, we performed sandwich ELISA. Briefly, 96-well Nunc MaxiSorp plates (Thermo Fisher Scientific) were coated overnight at 4°C with 10 μg/mL of either mAb Fab14 or FabE in carbonate buffer (pH 9.6) and then blocked for 2 hrs at 37°C using 5% BSA in PBST. Supernatant containing mutated capsids were added and incubated at room temperature for 1 hr. Wells were then washed three times with PBS with 0.05% Tween-20, then incubated with a rabbit anti-capsid polyclonal antibody for 1hr at room temperature. Donkey anti-human IgG (Fcγ fragment specific) antibody conjugated to horseradish peroxidase (HRPO) (Jackson ImmunoResearch Inc., West Grove, PA) was added and incubated for 1 hr at room temperature, then after washing 3,3’,5,5’-tetramethylbenzidine (TMB/ Thermo Fisher Scientific) was added and the optical density (OD) was measured at 450 nm with a Multiskan Ascent plate reader (Thermo Fisher Scientific). Binding of antibodies to WT CPV-2 capsids was run in parallel as positive controls.

### Virus neutralization assays

A neutralization assay was performed using CPV-2, where a virus stock that infected 13% of the mammalian cells in culture was mixed with 0.1, 1.0, and 10.0 μg/mL of the scFv-Fc and incubated for 1 hr at 37°C. Mixtures were then added to NLFK cells seeded at 2 × 10^4^ cells/cm^2^, and cultured for two days, then fixed and immunostained for infected cells using the mouse mAb against viral NS1 (48). The percentages of infected cells were determined for each condition and concentrations that neutralized ∼90% of infection were used in the selection studies.

### Experimental selection using mAbs and WT or mutant scFv-Fcs

Plasmid-derived WT CPV-2 virus was passaged ten times in NLFK cell cultures in the presence of either IgGs of mAb Fab14 or FabE, or engineered scFv-Fcs (WT and mutants). The selection was kept constant throughout the passages by adding fresh IgG or scFv at 0.3μg/mL for mAb or 1μg/mL for scFv-Fcs. Inoculums were prepared by mixing 63μL of virus (from the stock for the initial input or supernatant from the previous passage) with 7μL of antibody and incubated for 30 mins at 37 °C. The mixture was then added directly to cells and incubated for 1 hr at 37 °C, then the inoculum was removed, cells were washed, and media was added. After 3 days cells were frozen and thawed, and supernatant virus recovered. Controls (virus treated similarly but without antibodies or scFv-Fcs) were prepared and passaged in parallel.

### CVP-2 genome amplification, DNA libraries, and deep sequencing

Total DNA was isolated from clarified culture supernatants using the E.Z.N.A. Tissue DNA Kit as per the manufacturer’s instructions (cat. no D3396-01) and quantified via Qubit 4 Fluorometer (Thermo Fisher Scientific, cat. no. Q33238). The isolated DNA and the initial infectious plasmid were used as DNA template for polymerase chain reaction (PCR) using 30 cycles and the Q5 High Fidelity DNA Polymerase (New England BioLabs) under the recommended conditions by the manufacturer, amplifying the CPV-2 genome as two overlapping segments using the following primers: 5’-CCGTTACTGACATTCGCTTCTTG-3’ and 5’-GAACTGCTCCATCACT-CATTG-3’ (for NS1, ∼2.5 kb amplicon), and 5’-CATCCATCAACATCAAGACCAAC-3’ and 5’-CTTAACATATTCTAAGGGCAAACCAACCAA-3’ (for VP2, ∼2.5 kb amplicon). Each PCR product was purified with an 0.45 volume AMPure XP beads (Beckman Coulter), quantified via Qubit 4 fluorometer, and 0.5ng of each fragment mixed and used to construct barcoded sequencing libraries with the Nextera XT DNA Library Preparation Kit (Illumina, cat. no.FC-131-1096). Libraries were multiplexed and sequences were determined using Miseq 2 × 250 Illumina sequencing.

### Data analysis and bioinformatics

Raw sequencing reads were trimmed using BBDuk (https://jgi.doe.gov/data-and-tools/bbtools/bb-tools-user-guide/) to remove adaptors, PCR primers, and low-quality regions from reads. Reads were merged and mapped to a CPV-2 reference sequence (EU659116) using Geneious Prime v. 2019.0.4 (Kearse et al. 2012). Reads were error-corrected and normalized to 5000-fold coverage per site using BBNorm (https://jgi.doe.gov/data-and-tools/bbtools/bb-tools-user-guide/bbnorm-guide/). Reads for each sample were re-mapped to their consensus sequence and an additional filtering step (BAMUtil: Filt) (49) was performed to clip base miscalls near termini of reads. Coverage and site frequencies across the region of interest were determined using the PileupParam & ScanBamParam features in Rsamtools (50), and the minor variant frequencies called for each position. The accuracy and sensitivity of our approach to identifying rare sub-consensus single nucleotide variants in our viral samples have been previously tested on a control sequencing study as described by Voorhees *et al*. (28).

### Statistical analyses

All experiments were run in three independent replicates to address reproducibility, and data was analyzed using GraphPad Prism 9.3.1 (GraphPad Software, Inc., La Jolla, CA). Error bars represent the mean ± SEM, and statistical significance was defined at p≤0.05.

## RESULTS

### Capsid: antibody interface and the receptor-binding site

Previous work revealed the interactions between mAb Fab14 and FabE and CPV-2 capsids at high resolution (**Fig. 1A and B**) (33, 34). Here, we determined the selective pressure impose by those antibodies and defined the subsequent emergence of escape mutations in antibody and receptor binding sites (**Fig. 1A and B**). Importantly, this allowed us to define the variability of the epitopes recognized, to use antibody engineering to alter the Ab: capsid interactions, and address questions related to the natural evolution of CPV-2 in the face of the host humoral responses.

**Figure 1.**
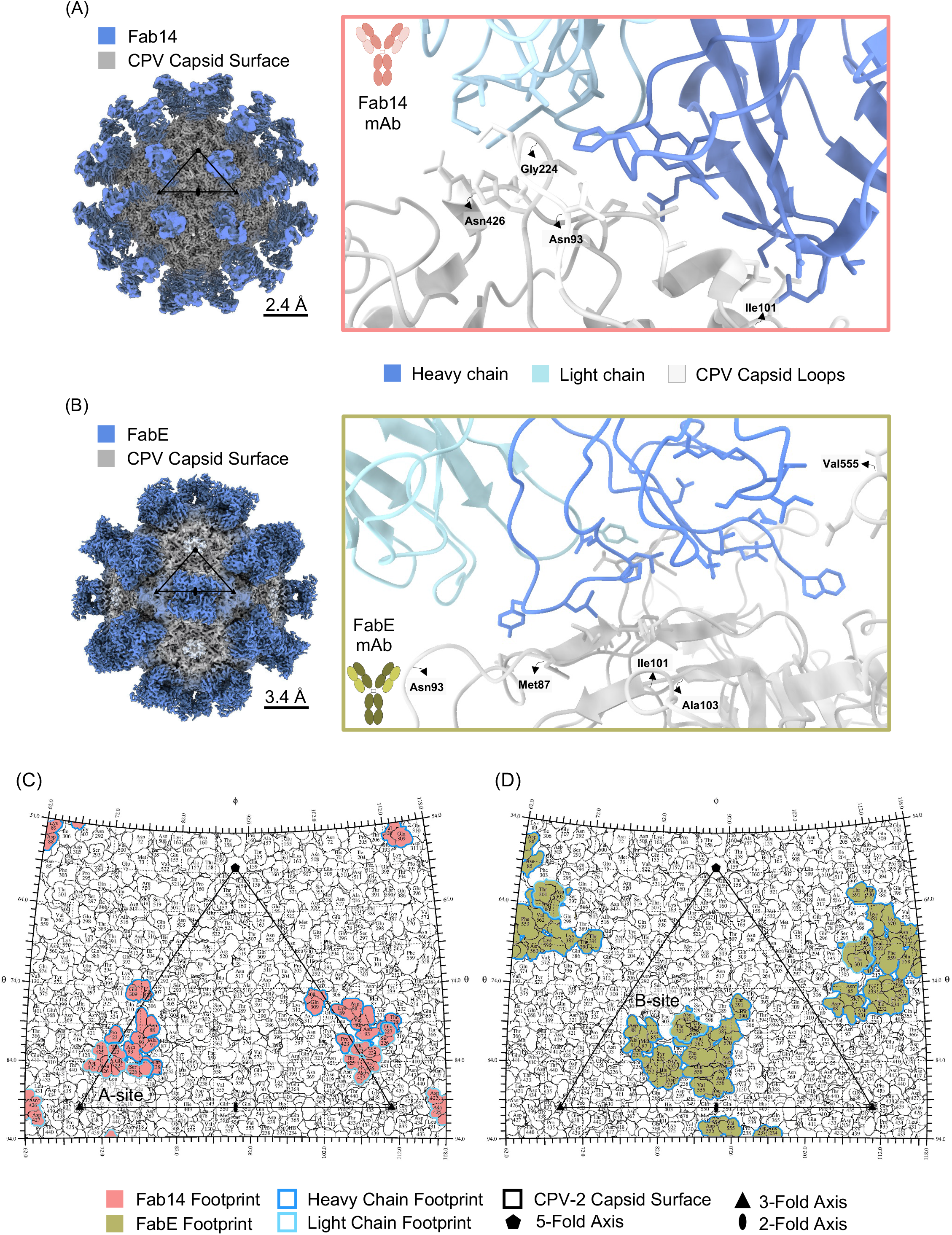
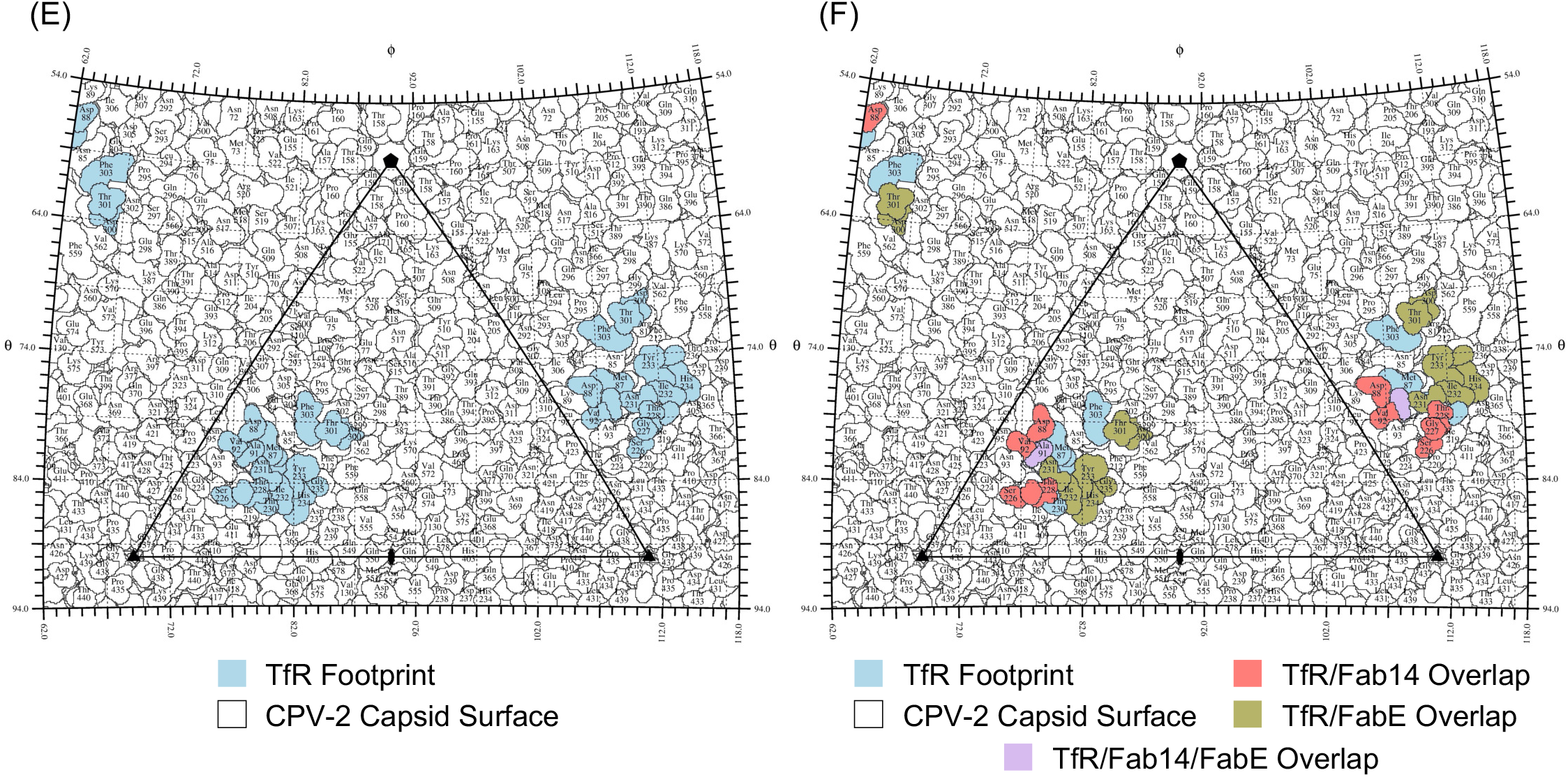
High-resolution cryoEM structures of the CPV-2-specific mAbs used in this study. (A, B) Reconstruction of A4E3 and B5A8 Fabs in complex with CPV capsids and their respective modeled mAb: capsid interface. (C,D) Roadmaps of A4E3 (A site, left) and B5A8 (B site, right) footprints on the CPV-2 capsid protein surface. Residues in the mAb footprints are lined based on their specific interaction with either the light or heavy chain antigen binding site. (E) Roadmap of the TfR apical domain footprint on the CPV-2 capsid surface. (F) Roadmap of the TfR apical domain footprint and the overlap with antigenic sites A and B. For all roadmaps, a single icosahedral subunit is denoted with a triangle and each axis of symmetry is represented by geometrical shapes.

Fab14 and FabE each have 60 potential binding sites on the CPV-2 capsid but bind to different positions on the threefold spike of the capsid: Fab14 binds close to the 3-fold axis (A-site), while FabE binds on the shoulder of that structure (B-site) (**Fig. 1C and D**). The capsid: antibody binding includes many different interactions, and the two binding sites overlap around VP2 residues Asp-88 and Ala-91. In both cases the heavy chain interacts with loops from three different VP1/2 subunits. Around 82% (14 of 17 residues in the receptor-binding footprint) of the TfR binding site overlap with the epitopes bound by the two antibodies (**Fig. 1E and F**).

### Deep sequencing of viruses and the effects of antibody selection

After passaging the viruses ten times in the presence of either antibody at concentrations that enforce significant constraint on viral growth, viral DNA was deep sequenced in a process that covered nearly the complete genome and lacking only the distal portions of the 5’- and 3’-termini (UTRs) (**Fig. 2A**). More than 5,000-fold coverage per site was achieved after read processing and normalization. A short region (∼150 nucleotides in length) with a poly(A)- and G-rich sequence region close to the start of the VP2 gene, as well as sequences close to the 5’-UTR, did not sequence well for either viral DNA samples or the control plasmid. These “low coverage regions” [LCR] in **Fig. 2B** did not affect consensus calls but did impact the calling of SNVs frequencies and were therefore excluded from sequence diversity measurements.

**Figure 2.**
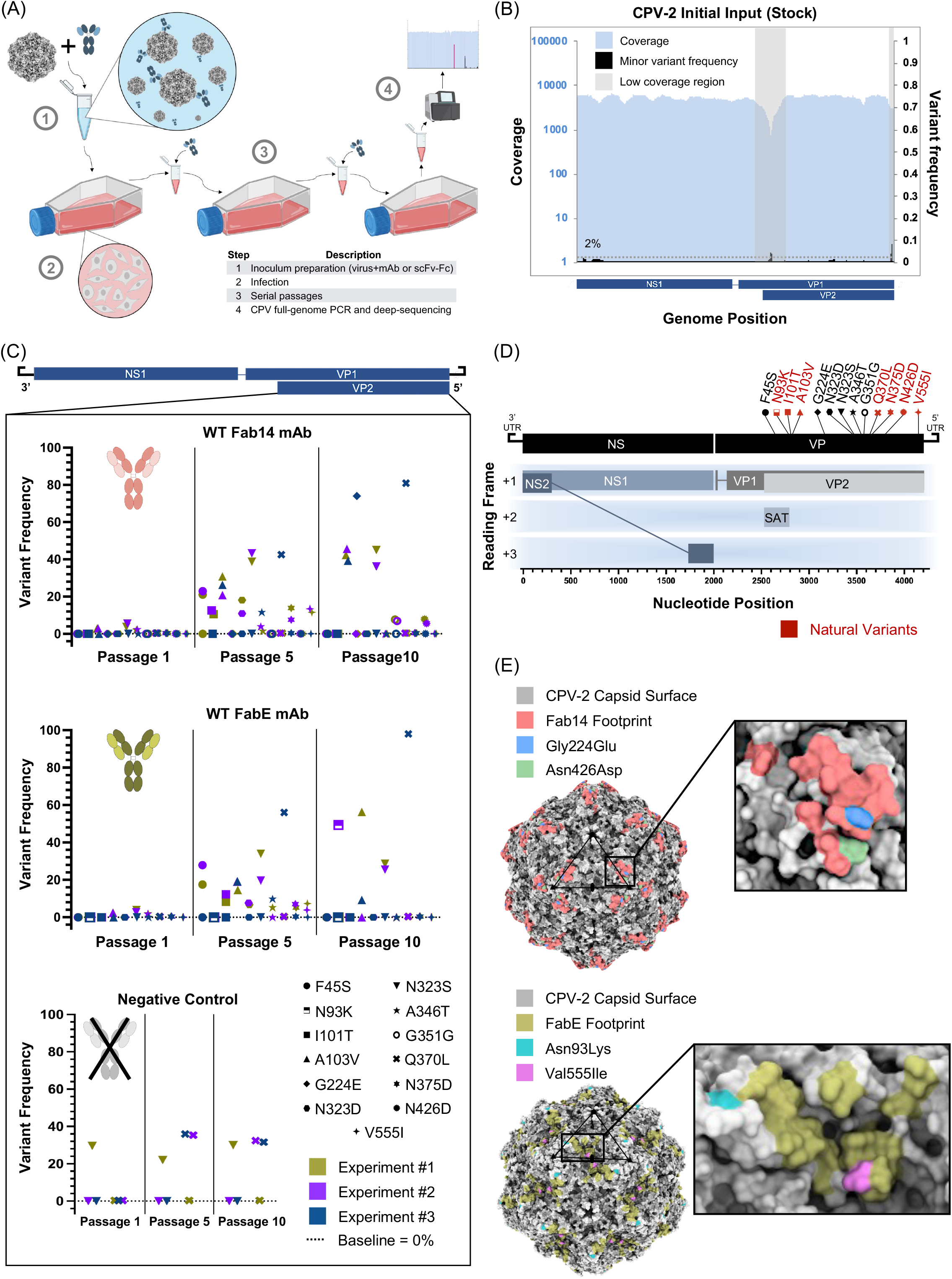
*In-vitro* antibody selection on CPV-2 viruses with mAbs. (A) Schematic of the experimental approach to select CPV-2 virus in the presence of A4E3 and B5A8 mAbs. The numbers indicate each step in the experiment and the respective description is included in the legend. (B) The genetic background of the CPV-2 input virus that was used in all the experimental passages. The <low coverage regions= (LCR) are denoted with gray boxes in relation to the parvovirus genome. (C) Summary of the changes in single-nucleotide variants (SNVs) frequencies in the VP2 gene throughout the antibody selection experiment for each mAb and the negative control group. Each shape represents a different SNV, and the colors represent each independent replicate (n=3). The baseline is denoted at frequency of 0% (dotted line). (D) Survey of the antibody selected mutations in relation to the position in the CPV-2 ssDNA genome. Variants found in nature are denoted in red. (E) Localization of the antibody selected mutations exposed on the surface of the CPV-2 capsid that fall into the A4E3 and B5A8 footprints (reconstructed high-resolution CPV-2 capsid from Lee et al., 2019). The capsid reconstruction was modeled in Chimera X.

This analysis can consistently detect SNVs at low proportions (1% or less) across the CPV-2 genome. Accordingly, we set a cutoff of 1% for SNV detection (28). The input viruses showed few SNVs above 1%, and all were less than 2% (**Fig. 2B**). In contrast, all viruses incubated with mAb Fab14 or FabE showed SNVs at higher levels, with almost all being non-synonymous and within the VP2 gene (**Fig. 2C**). After the first passage three nonsynonymous SNVs arose in the VP2 gene at low frequencies (<10%) in viruses incubated with either of the mAbs: VP2 A103V, N323S, and A346T (**Fig. 2C**), and by passage five we also detected changes in positions VP2 F45S, I101T, A103V, N323D, N323S, A346T, Q370L, N375D, and V554I (**Fig. 2C**). By passage ten, SNVs were detected for each selected viral population. MAb Fab14-selected viruses showed VP2 mutations A103V, G224E, N323S, and Q370L at frequencies of ∼40% or higher, while VP2 G351G and N426D were present at ∼8% in frequency. MAb FabE-selected viruses showed mutations VP2 N93K, A103V, N323S, and Q370L at 20% frequency or higher. Virus passaged ten times without antibody also showed increased frequencies of SNVs VP2 N323S and Q370L (<40% frequency, **Fig. 2C**), indicating that they were selected by a non-antibody component of the culture system. Strikingly, all SNVs fell within the CPV-2 VP2 gene (**Fig. 2D**) and almost all were nonsynonymous changes with the exception of VP2 G351G. Of the antibody-selected SNVs detected, 54% have been found in one or more natural circulating CPV-2 viruses (SNVs are highlighted in red in **Fig. 2D**) (28). Other synonymous SNVs in the parvoviral genome remained below 2% of frequency and were not analyzed further.

We mapped the SNVs selected onto the viral capsid structures (**Figs. 2C and D**), which showed that VP2 G224E and N426D fall directly within the Fab14 footprint, whereas VP2 V555I fall directly into the FabE footprint and VP2 N93K is near to that region (**Fig. 2E**). All other SNVs - with the exception of VP2 N375D - are either close to or underneath both antibody binding sites. None of the antibody selected SNVs fell directly into the footprint of the apical domain of TfR (TfR footprint shown in **Fig. 1E** and TfR: mAb footprint overlapping in **Fig. 1F**).

### Antibody-selected mutations in the viral capsid reduce antibody binding

To examine the effect of antibody-selected mutations on mAb interactions, we mutated two surface-exposed residues (VP2 G224E and N426D) and one internal residue (VP2 I101T) in the CPV capsid structure and screen for variations in antibody binding. All tested capsid mutants showed reduced ELISA binding to both mAbs compared to the WT (**Figs. 3A and B**), and VP2 N426D had the greater effect on binding of for both mAbs, and many mutations selected by one of the mAbs in cell culture also affected the overall binding of the other mAb.

**Figure 3.**
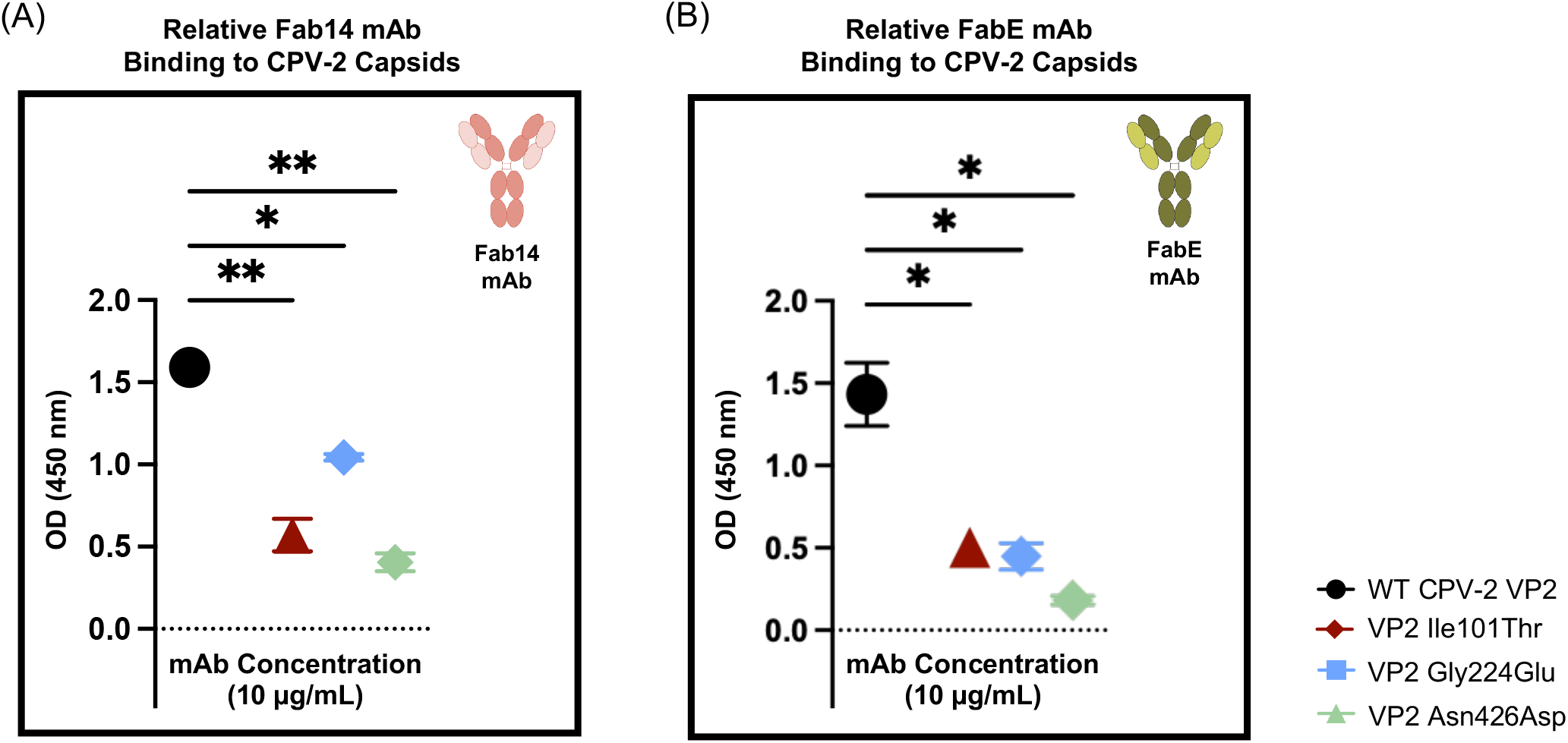
Relative binding of both mAbs to mutated CPV-2 capsids with antibody-selected mutations. The effect of antibody-selected mutations in the CPV-2 capsids (VP2) on binding to mAbs determined by ELISA. (A) Relative binding of mutated CPV-2 capsids (VP2) to A4E3 mAb. (B) Relative binding of mutated CPV-2 capsids (VP2) to B5A8 mAb. The quantification was normalized to the background signal in absence of mAbs. Intact CPV-2 capsids (WT CPV-2) were used as positive control. All experiments were conducted in three independent replicates (n=3), and all mAbs were tested at 10 μg/mL. Error bars show mean ± standard mean error. Statistics were calculated using ANOVA. *, *P* < 0.05; **, *P* < 0.002.

### Effects of antibodies with engineered antigen-binding sites

To better understand how the specific antigen binding sites of the antibodies influence the mutational selection patterns in CPV-2, we engineered the CDRs of six antibodies as scFv-Fcs (**Fig. 4A**). Four were single-amino acid mutants (Lc N30D and Hc T30K in Fab14 scFv-Fcs, and Hc M152D and Hc P202E in FabE scFv-Fcs) and two were double-mutants (N30D/T30K in the Lc and Hc, respectively, of Fab14 scFv-Fcs; and M152D/P202E, both in the Hc of FabE scFv-Fcs) as shown in **Fig. 4B and C**. We also prepared the WT scFv-Fc of each antibody.

**Figure 4.**
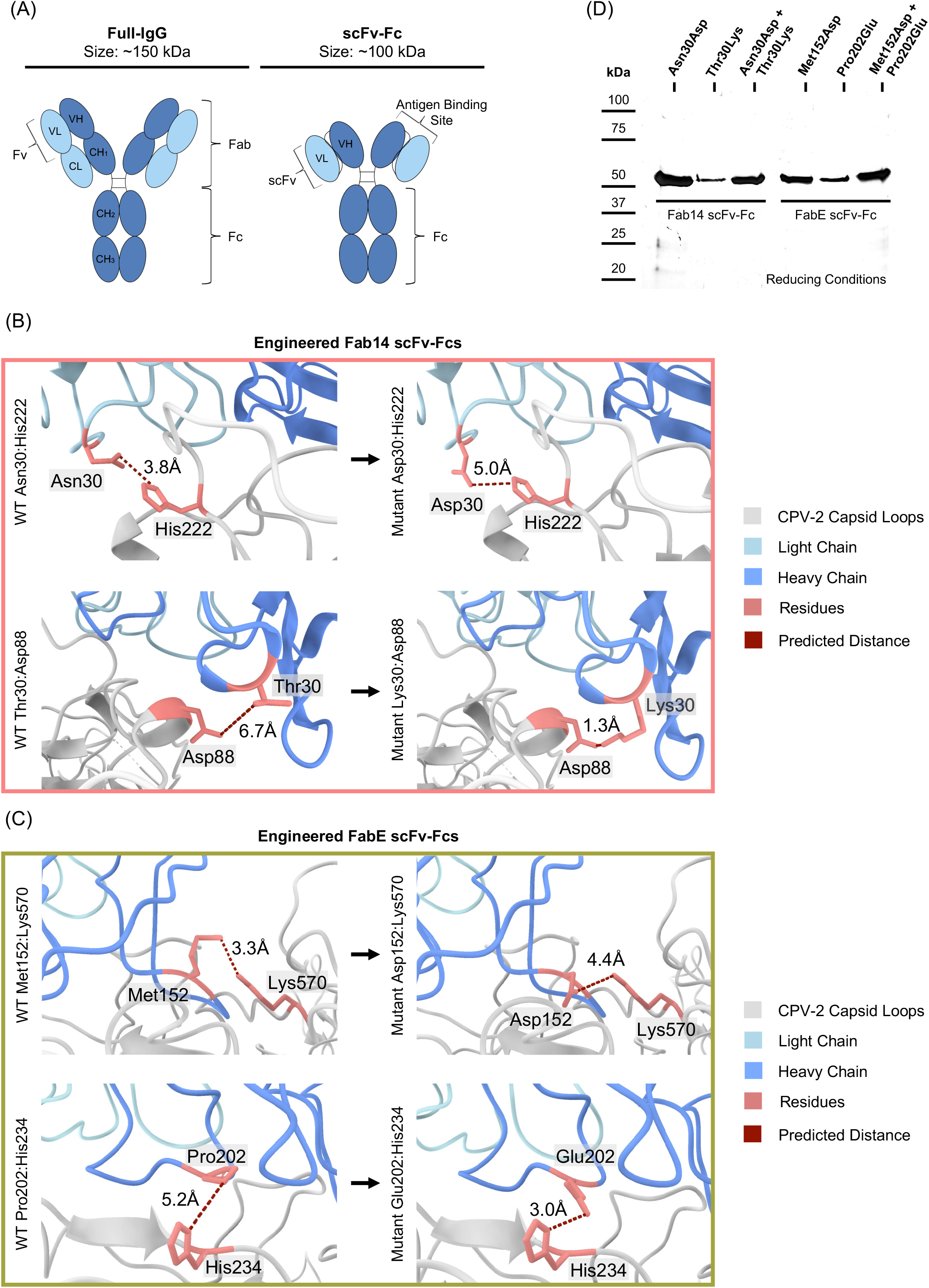
Engineered antigen-binding sites of CPV-2 specific mAbs as scFv-Fcs. (A) Side-by-side comparison of a full-IgG with an scFv-Fc and the structural features present in each molecule. (B, C) Reconstructed models for each engineered scFv-Fc antigen-binding site showing the predicted residue conformation (in salmon) and the relative atomic distance between fully-charge residues present in the antigen-binding site complementarity-determining region and the CPV-2 capsid loop (atomic distance in red and CPV-2 capsid loops in gray). Light chains are denoted in light blue and heavy chains are denoted in darker blue. Engineered A4E3 scFv-Fcs are in panel B and engineered B5A8 scFv-Fcs are in panel C. (D) Silver staining showing FPLC-purified engineered scFv-Fcs. The antibody: capsid interaction models were performed in Chimera X.

Each protein was expressed from baculovirus vectors in insect cells, and purity was assessed using silver-stain in reducing conditions (**Fig. 4D**). ELISA showed these scFv-Fcs had various binding levels, with some showing the same binding affinity as WT, while others showed reduced binding (**Figs. 5A and D**). Engineered scFv-Fcs showing similar binding ELISA compared to their WT counterparts also showed similar neutralization to the WT scFv-Fcs, while scFv-Fcs with lower ELISA binding also neutralized infectivity to lower levels compared to their WT counterparts (**Figs. 5B, C, E, and F**).

**Figure 5.**
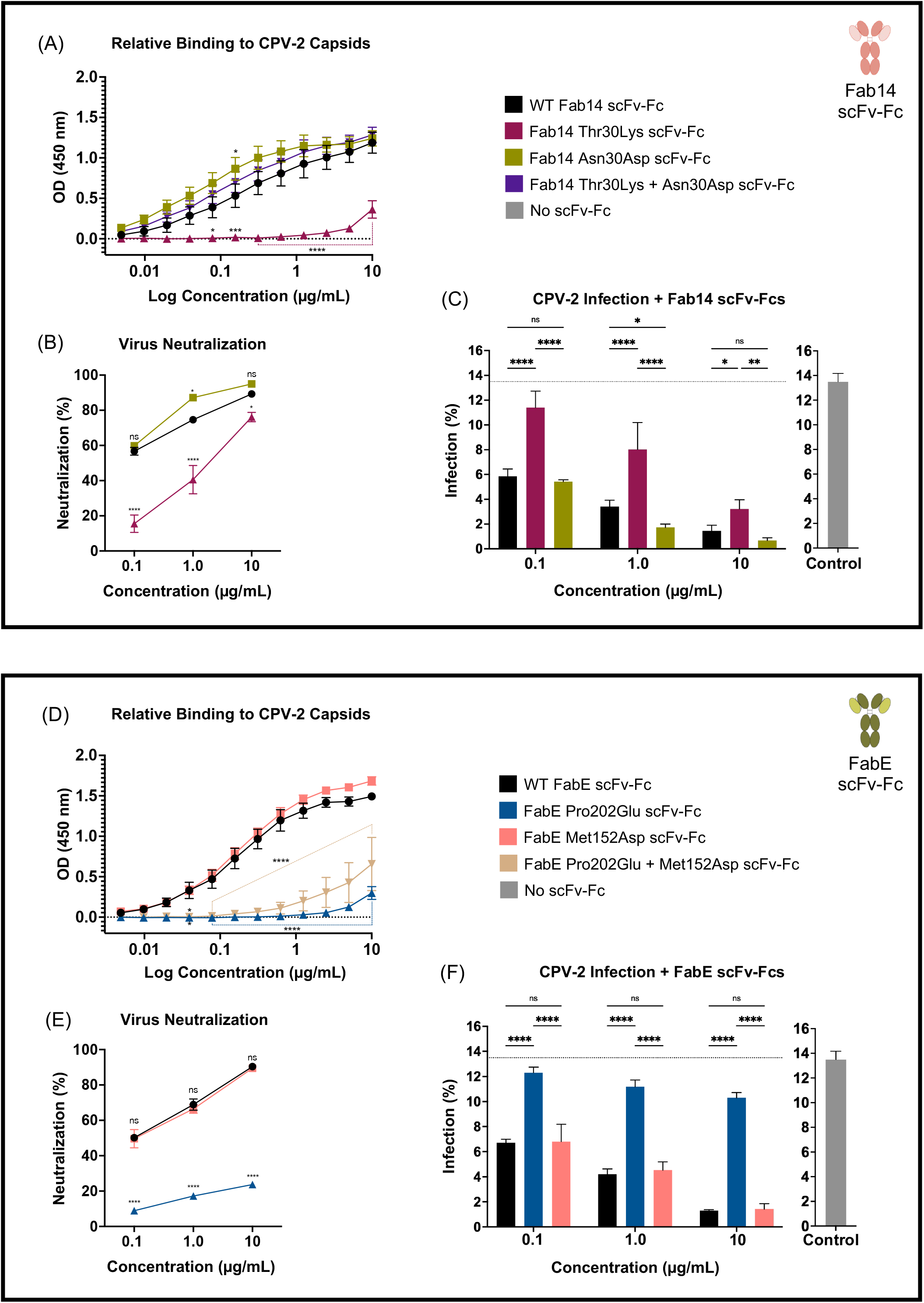
Relative binding CPV-2 capsid binding, virus neutralization, and infectivity assays to characterize engineered scFv-Fcs. To characterize all engineered scFv-Fcs, we tested for their relative binding to CPV-2 capsids, and their effect on CPV-2 neutralization and infection *in-vitro*. Panels A, B, and C correspond to experiments using A4E3 scFv-Fcs and panels D, E, and F correspond to experiment using B5A8 scFv-Fcs. (A, D) Relative binding of each engineered scFv-Fc to CPV-2 capsids tested by ELISA. (B, E) Quantification of neutralization of CPV-2 capsids upon infection *in-vitro* in the presence of two engineered scFv-Fcs (a WT-like binder and a poor binder). (C, F) Quantification of CPV-2 infected cells after incubation with each engineered scFv-Fc. The quantification was normalized to the maximum number of CPV-2 infected cells in cell culture in the absence of scFv-Fcs. All experiments were conducted in three independent replicates (n=3), and all scFv-Fcs were tested at the same three different experimental concentrations. Error bars show mean ± standard mean error. Statistics were calculated using ANOVA. *, *P* < 0.05; **, *P* < 0.002; ***, *P* = 0.0001; ****, *P* < 0.0001.

For BLI measurements of binding, the scFv-Fcs were bound to protein A biosensors then incubated with CPV-2 capsids (**Fig. 6A**). WT Fab14 and FabE scFv-Fcs showed high affinities, with both fast association rates and very low dissociation rates (**Figs. 6B and C**). Engineered scFv-Fcs Fab14 Lc N30D and FabE Hc M152D that showed similar binding to their WT counterparts in ELISA (**Figs. 5A and D**) showed decreased association rates by BLI (**Figs. 6 B and C**). Although scFv-Fc FabE Hc M152D showed very low dissociation rates similar to its WT counterpart, scFv-Fc Fab14 Lc N30D showed high off-rates (**Figs. 6B and C**). The scFv-Fc Fab14 Hc T30K or FabE Hc P202E did not bind CPV-2 capsids in BLI (**Figs. 6B and C**).

**Figure 6.**
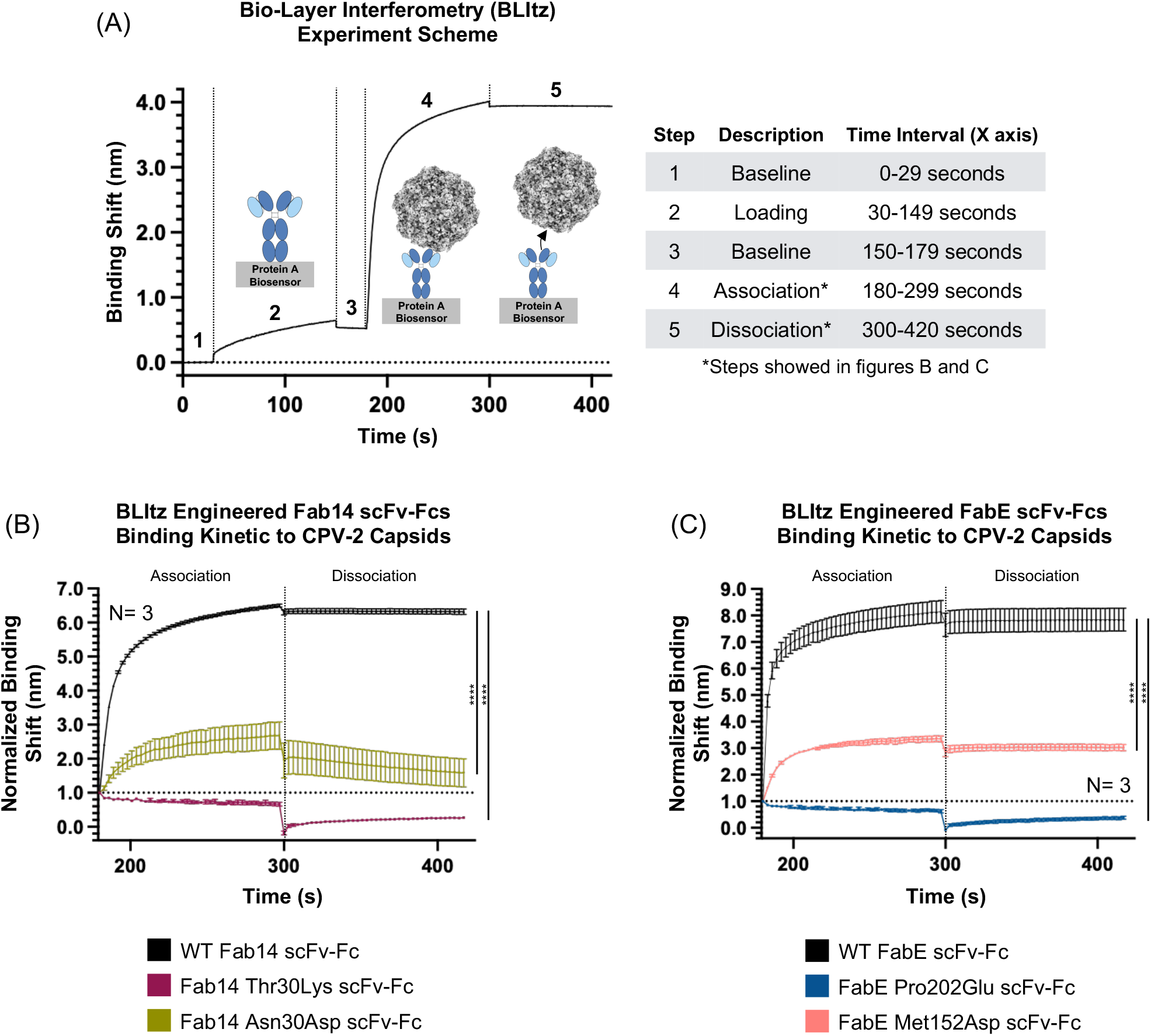
Binding kinetics of CPV-2 capsids to engineered scFv-Fcs. (A) A sample bio-layer interferometry experiment. Protein A biosensors were washed (step 1), incubated with scFv-Fcs (step 2), washed again (step 3), incubated with CPV-2 capsids to measure scFv-Fc/capsid association (step 4), and washed to measure scFv-Fc/capsid dissociation (step 5). Results are measured as the change in the wavelength (in nm) over time and normalized to the baseline read prior to the loading step. (B) Capsid binding kinetics to A4E3 scFv-Fcs. (C) Capsid binding kinetics to B5A8 scFv-Fcs. All experiments were conducted in three independent replicates (n=3). Error bars show mean ± standard mean error. Statistics were calculated for the 180-s through the 420-s time points using ANOVA. ****, *P* < 0.0001.

### Antibody selection using engineered antibodies

We used the WT and some of the engineered scFv-Fc (**Fig. 4B and C**) for selection in cell culture (**Fig. 2A**) and found very similar results to those seen after mAb selection. The same initial virus stock was used for the selection, and we again found only a small number of SNVs above 1%, and all were less than 2% with the exception of the LCRs (**Fig. 7A**). All viruses incubated with scFv-Fcs, including the WT Fab14 and FabE scFv-Fcs and their mutated forms, showed SNVs at higher levels (**Fig. 7B and C**). After the first passage, we detected three nonsynonymous SNVs in the VP2 gene in viruses incubated with all scFv-Fc forms: VP2 A103V, A346T, and N323S (**Fig. 7C**). Those were the same as after selection with mAbs but were at higher frequencies (around 10% to 40%), with only a few other nonsynonymous VP2 SNVs at lower frequencies (<10%). By passage five we also detected changes in positions VP2 M87L, I101T, T194A, V316I, N375D, R377K, V423I, N426D, and V555I, all were present at low frequencies (<10%, **Fig. 7C**). We also observed an increase in frequency for SNV VP2 A103V (around 70% to 100%), while VP2 A346T and N323S remained at lower frequencies (10% to 40%, **Fig. 7C**). Viruses passaged five times without antibody again showed increased frequencies of SNVs VP2 N323S and N375D (<40% frequency, **Fig. 7C**). In these studies, we did not see a change of VP2 residue 370L. However, this again indicated selection by a non-antibody component of the system. Interestingly, 7 out of the 12 SNVs found after scFv-Fc selection were also selected with mAbs, including VP2 I101T, A103V, N323S, A346T, N375D, and V555I. As was seen for selection by mAbs, all antibody-selected SNVs fell within the CPV-2 VP2 gene (**Fig. 7B**), and all were nonsynonymous (**Fig. 2D**). As seen before, 50% of the changes have been found in one or more natural circulating CPV-2 viruses as reported in our previous studies (these SNVs are highlighted in red in **Fig. 7B**) (28).

**Figure 7.**
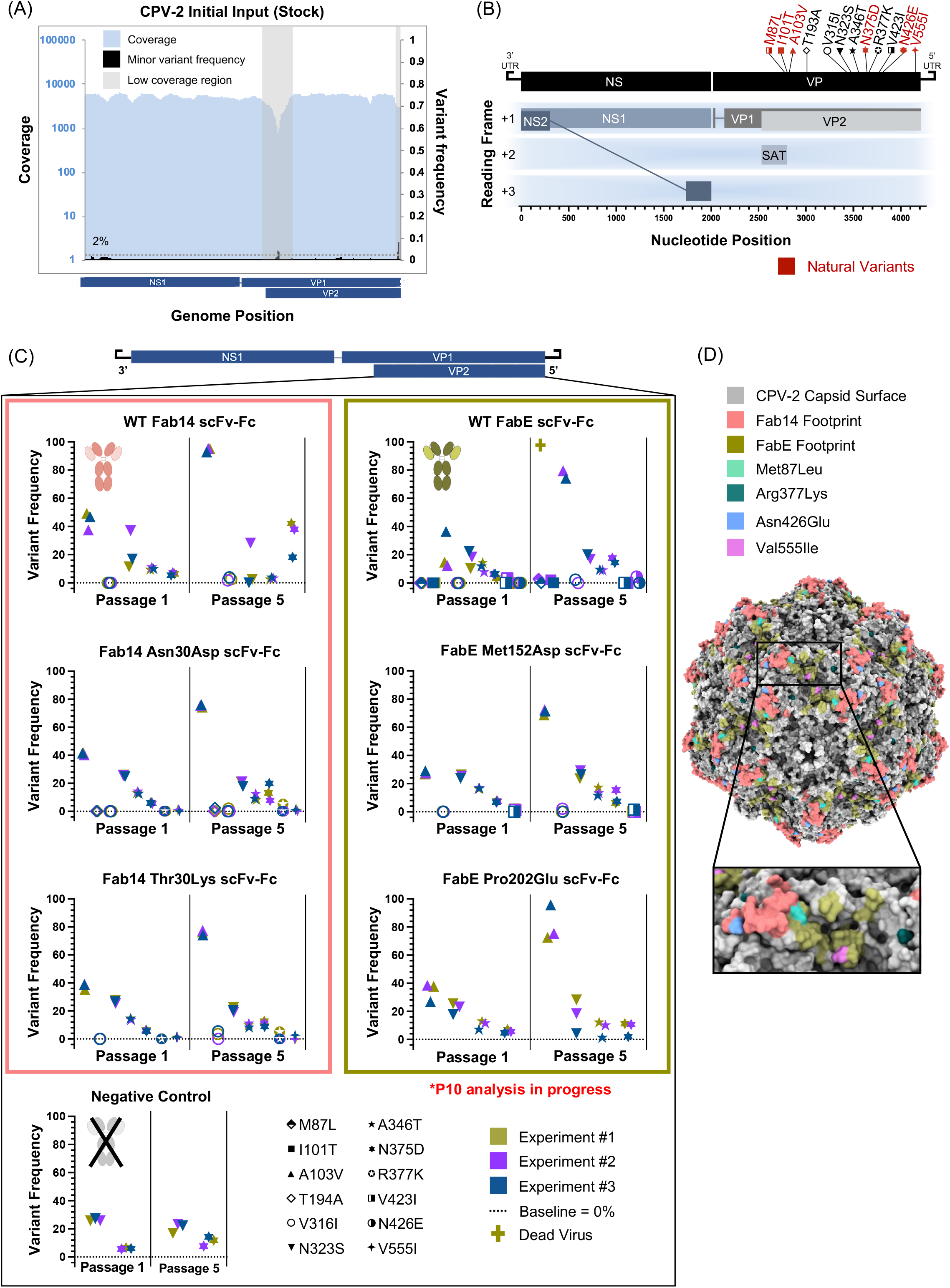
*In-vitro* antibody selection on CPV-2 viruses with engineered scFv-Fcs. (A) The genetic background of the CPV-2 input virus was used in all the experimental passages. The <low coverage regions= (LCR) are denoted with gray boxes in relation to the parvovirus genome. (B) Survey of the antibody selected mutations in relation to the position in the CPV-2 ssDNA genome. Variants found in nature are denoted in red. (B) Survey of the changes in the frequency of SNVs in CPV-2 genomes throughout the antibody selection experiments using WT and engineered A4E3 scFv-Fcs, N30D and T30K (left panels in salmon colored box); WT and engineered B5A8 scFv-Fcs, M152D and P202E (right panels in green colored box); and the negative control group. Each shape represents a different SNV, and the colors represent each independent replicate (n=3). The baseline is denoted at frequency of 0% (dotted line). (D) Localization of the antibody selected mutations exposed on the surface of the CPV-2 capsid that fall into the A4E3 and B5A8 footprints (reconstructed high-resolution CPV-2 capsid from Lee et al., 2019). The capsid reconstruction was modeled in Chimera X.

Mapping the SNVs that arose after scFv-Fc selection showed that SNV VP2 N426D fall directly within the Fab14 footprint, while SNVs VP2 M87L and V555I fell directly in the FabE footprint (**Fig. 7D**). All other SNVs – with the exception of VP2 N375D - are either close to or underneath both antibody binding sites. None of the antibody selected SNVs fell directly into the footprint of the apical domain of TfR, with exception of SNV VP2 M87L, which did not exceed a frequency lower than 5% (TfR footprint shown in **Fig. 1E** and TfR: mAb footprint overlapping in **Fig. 1F**).

## DISCUSSION

We conducted an experimental evolutionary analysis of a DNA virus undergoing selection by two different neutralizing mAbs and characterize the viral sequence and structural variation that arose during passage in cell culture. The variation and evolution of antigenic structures are intrinsic properties of both RNA and DNA viruses, and selection of escape mutations by mAb has often been used to define antigenic epitopes and to understand the nature of immune escape (35–37). However, the details of how these mutations arise and are selected, as well as the relationships between antibody selection and receptor binding, are still not well understood despite the overlaps of their binding sites in many viruses. This study therefore has implications for understanding the molecular properties and selection of viral epitopes and how those relate to receptor binding sites, such as those seen in other viral systems including SARS-CoV-2, HIV-1, influenza, Nipah, some adeno-associated viruses, and human picornaviruses (6, 51–57).

### Mutations in parvoviruses and the role of selection

The CPV strain studied here has undergone a well-defined natural evolution that involves both antigenic and receptor binding sites (26, 28, 58, 59). The capsid itself is simple, being comprised of 60 repeated structural subunits which each has only a relatively small, exposed surface area. The virus stock was isolated from a plasmid and grown for a few passages, so that little preexisting variation was present in the virus we started with. However, after passaging in the presence of each of the two antibodies tested here, mutations were rapidly selected in the virus, confirming that parvovirus DNA replication results in the generation of mutations that remain at very low levels without the exertion of a selection pressure. The antibodies used here only selected mutations within the VP2 gene, and all except one resulted in non-synonymous changes. No mutations that reached a frequency of >2% were seen in the NS1 gene or in the sequence encoding the VP1 unique region buried inside the capsid. Hence, these results therefore mirror those seen for viral samples recovered from infected animals, where only a small number of subconsensus variants were observed in the full genome deep sequences (28).

Two mutations arose in the control viruses passaged in cells without antibodies, and these mutations were also on the capsid surface, suggesting that a component of the cells or the culture system was selective under the conditions of these experiments. NLFK feline cells have been widely used in previous studies, and have occasionally been seen to select for specific mutations variations in the virus, including the loss of sialic acid binding likely due to the expression of N-glycolyl neuraminic acid (Neu5Gc) in feline cells (15).

### Properties of antibody selected mutations

After antibody selection, altered residues were all exposed on or immediately under the capsid surface. Most mutations were within or one residue away from the known binding site of the selecting antibody, and several were within a region within the footprints of both antibodies. Our Illumina sequencing produced sequence reads of 250 bp, sufficient for us to confirm that closely spaced mutations were not linked (i.e., VP2 N93K, I101T and A103V; VP2 N323D/S, A346T, G351G, Q370L, and N375D; and VP2 N426D and V55I). Most mutations that were further apart also appeared not to be linked as they were selected and changed in proportion separately. We were also able to confirm that some of the antibody-selected mutations resulted in viruses with reduce binding affinity to both mAbs confirming that those are escape mutants.

In addition to the wildtype antibodies, we examined the effects of the specific antibody structures and interactions by altering positions predicted from the structures of their complexes with the capsids. The wildtype scFv-Fcs bound with similar kinetics to the original IgG (60), and some of the mutated scFv-Fcs showed reduced binding compared to the wildtype. These were used to select the virus. Despite the structural changes and reduced affinity of those scFv, all mutations selected by the mutated scFv-Fcs were within the same spectrum of changes as seen for the wildtype scFvs or the original IgGs.

### Overlap with the receptor binding site, including the canine TfR glycan

The virus binds to the TfRs of various hosts to enter and infect cells. The binding site of the blackbacked jackal TfR has been determined by cryoEM, showing the complex between the TfR protein structure and the capsid (18). However, the canine TfR contains an additional glycan attached to VP2 Asn 384 of that structure, and that glycan also interacts with the capsid (27). We therefore examined both the protein: protein interactions, as well as the capsid residues that were close to the glycan interacting structure. From all antibody-selected mutations, only a mutation of VP2 residue 224 (G224E) was located near the TfR footprint and therefore likely to influence the host receptor binding, in addition to their effects on the antibody binding. This restricted number of mutations in TfR-interacting residues may allow parvoviruses to acquire antibody-evading mutations that avoid detrimental effects on the TfR: capsid interaction required for cellular infection.

### Connections to the natural evolution of CPV and FPV

Selection with these rodent mAb resulted in the isolation of many mutations that were the same as those identified a full-genome deep-sequence analysis of the CPV genomes in natural samples collected between 1978 and 2019 (those SNVs are highlighted in in **Fig. 2D**) (28). That many of these residues were multiply selected during our i*n vitro* studies indicates that there are only a small number of mutations that introduce antigenic escape while retaining efficient viral replication. These were likely due to both intrinsic structure and functional features of the capsid, as well as the need to maintain functional TfR binding. In the case of dog cells these binding includes both the protein and glycan structures.

## ACKNOWLEDGEMENTS

Antibody: capsid structures and molecular analyses were performed with UCSF Chimera X, developed by the Resource for Biocomputing, Visualization, and Informatics at the University of California, San Francisco, with support from National Institutes of Health R01-GM129325 and the Office of Cyber Infrastructure and Computational Biology, National Institute of Allergy, and Infectious Diseases.

## SUPPORT

This work was supported by the National Institutes of Health grants R01-AI092571 and R01-GM080533 to Colin R. Parrish, the NIH-Diversity Supplement 3R01AI092571-08W1 to Colin R. Parrish and Robert López-Astacio, and the Daversa Family Scholarship to Robert López-Astacio.

